# Lack of antiviral activity of probenecid in Vero E6 cells and Syrian golden hamsters: a need for better understanding of inter-lab differences in preclinical assays

**DOI:** 10.1101/2022.03.03.482788

**Authors:** Helen Box, Shaun H Pennington, Edyta Kijak, Lee Tatham, Claire H Caygill, Rose C Lopeman, Laura N Jeffreys, Joanne Herriott, Joanne Sharp, Megan Neary, Anthony Valentijn, Henry Pertinez, Paul Curley, Usman Arshad, Rajith KR Rajoli, Steve Rannard, James P. Stewart, Giancarlo A Biagini, Andrew Owen

## Abstract

Antiviral interventions are urgently required to support vaccination programmes and reduce the global burden of COVID-19. Prior to initiation of large-scale clinical trials, robust preclinical data in support of candidate plausibility are required. The speed at which preclinical models have been developed during the pandemic are unprecedented but there is a vital need for standardisation and assessment of the Critical Quality Attributes. This work provides cross-validation for the recent report demonstrating potent antiviral activity of probenecid against SARS-CoV-2 in preclinical models (1). Vero E6 cells were pre-incubated with probenecid, across a 7-point concentration range, or control media for 2 hours before infection with SARS-CoV-2 (SARS-CoV-2/Human/Liverpool/REMRQ0001/2020, Pango B; MOI 0.05). Probenecid or control media was then reapplied and plates incubated for 48 hours. Cells were fixed with 4% v/v paraformaldehyde, stained with crystal violet and cytopathic activity quantified by spectrophotometry at 590 nm. Syrian golden hamsters (n=5 per group) were intranasally inoculated with virus (SARS-CoV-2 Delta variant B.1.617.2; 103 PFU/hamster) for 24 hours prior to treatment. Hamsters were treated with probenecid or vehicle for 4 doses. Hamsters were ethically euthanised before quantification of total and sub-genomic pulmonary viral RNAs. No inhibition of cytopathic activity was observed for probenecid at any concentration in Vero E6 cells. Furthermore, no reduction in either total or sub-genomic RNA was observed in terminal lung samples from hamsters on day 3 (P > 0.05). Body weight of uninfected hamsters remained stable throughout the course of the experiment whereas both probenecid- (6 - 9% over 3 days) and vehicle-treated (5 - 10% over 3 days) infected hamsters lost body weight which was comparable in magnitude (P > 0.5). The presented data do not support probenecid as a SARS-CoV-2 antiviral. These data do not support use of probenecid in COVID-19 and further analysis is required prior to initiation of clinical trials to investigate the potential utility of this drug.

## Introduction

A concerted global effort over the first two years of the pandemic has sought to rapidly evaluate antiviral drug candidates to complement the highly successful, rigorously validated but still highly unequitable vaccination programmes. Many clinical trials have been and continue to be focussed upon investigating putative antiviral drugs that have been repurposed either after approval for another indication (e.g. hydroxychloroquine, lopinavir, ivermectin (1–7)) or in various stages of development for other indications (e.g. remdesivir, molnupiravir, nirmatrelvir (8–12)). The speed at which drugs can be brought forward under the urgency of a pandemic is a significant advantage. However, this strategy is prone to failure in the absence of robustly conducted and validated preclinical data.

Clinical trials incur significant costs and place additional burden on the healthcare staff needed for their conduct (13, 14). It is therefore critical that only candidates that can be robustly justified are studied. Candidates should only be considered worthy of investigation if (i) the mechanism of action is plausible and supports the intended use, (ii) the pharmacokinetics at the proposed dose support that antiviral activity can be achieved in the target population, (iii) reproducible preclinical data are available to demonstrate activity in preclinical models, (iv) acceptable safety in the target population can be justified. Although the safety and pharmacokinetics of drugs repurposed after approval for another indication is usually well understood, the putative impact of the disease should also be considered, particularly when known adverse drug effects may overlap with disease symptomology. Caution is also required since pharmacokinetics in COVID-19 patients can differ to that in patients with the primary indication, as was clearly demonstrated for lopinavir where drug exposure was considerably higher in COVID-19 patient populations than was observed in HIV patients (15). Under the urgency of pandemic conditions in particular, gaps in knowledge exacerbate risks and a priori expectations about performance cannot be guaranteed when drugs are applied to treat a condition different to that for which the drug was originally developed. Robust preclinical proof of efficacy data should be a prerequisite for selection of candidates for clinical trials.

During the pandemic, preclinical models of SARS-CoV-2 infection for the assessment of drug candidates have been developed at unprecedented speed and data to either support or refute candidacy of repurposing opportunities has been forthcoming (16–20). However, cross-validation of preclinical supporting evidence is needed to support a recommendation to progress a drug from preclinical testing to clinical trials.

Probenecid is a gout treatment that has previously been studied for antiviral drug repurposing in influenza (21). Recently, Murray et al. described highly promising preclinical activity of probenecid against SARS-CoV-2 with in vitro IC50 values as low as 0.00001 μM and a 3-4 log drop in viral load in SARS-CoV-2-infected Syrian golden hamsters (22). At the time of writing, a clear mechanism of antiviral action of probenecid for SARS-CoV-2 has not been empirically evidenced. However, the low cost, favourable safety profile and wide availability of the drug would advance the implementation of this treatment if antiviral activity was confirmed. Accordingly, the present study sought to replicate prior findings to provide additional data to support decision-making for clinical evaluation of probenecid.

## Methods

### Materials

Phosphate buffered saline (PBS) was purchased from Merck. Male Syrian golden hamsters were purchased from Janvier Labs. 1 mL Amies Regular flocked swab were purchased from Appleton Woods. GoTaq^®^ Probe 1-Step RT-qPCR System was purchased from Promega. SARS-CoV-2 (2019- nCoV) CDC qPCR Probe Assay, CDC RUO 2019-nCoV_N_Positive Control and the SARS-CoV-2 E SgRNA were purchased from IDT. TRIzol reagent, GlycoBlue™, Phasemaker™ tubes, Nanodrop and TURBO DNA-free™ kit were purchased from ThermoFisher. A bead mill homogeniser was purchased from Fisher Scientific. Precellys CKmix lysing tubes were purchased from Bertin Instruments. A Chromo4™ Real-Time PCR Detector purchased from Bio-Rad. Transmission cages were purchased from Techniplast UK Ltd.

### Viral isolates

A UK strain of SARS-CoV-2 (hCoV-2/human/Liverpool/REMRQ0001/2020), Pango lineage B, was cultured from a nasopharyngeal swab from a patient(23). The sequence was submitted to Genbank, accession number MW041156.

The B.1.617.2 (Delta variant) hCoV-19/England/SHEF-10E8F3B/2021 (GISAID accession number EPI_ISL_1731019), was kindly provided by Prof. Wendy Barclay, Imperial College London, London, UK through the Genotype-to-Phenotype National Virology Consortium (G2P-UK). Sequencing confirmed it contained the spike protein mutations T19R, K77R, G142D, Δ156-157/R158G, A222V, L452R, T478K, D614G, P681R, D950N.

The titres of all isolates were confirmed on Vero E6 cells and the sequences of all stocks confirmed.

### In vitro Vero E6 cell assay

7-point concentration-effect analysis was performed with probenecid in 96-well plates using VERO E6 cells. Cells were preincubated with probenecid or remdesivier (control) at 25.00 μM, 8.33 μM, 2.78 μM, 0.93 μM, 0.31 μM, 0.10 μM and 0.03 μM, or control media at 37°C with 5% CO2 for 2 hours. Preincubation media was removed and replaced with 50 μL minimal media containing SARS-CoV-2 (MOI of 0.05), 100 μL 2 × semi-solid media and then 50 μL minimal media containing probenecid, remdesivier (control) or control media, as appropriate. The plates were then incubated at 37°C with 5% CO2. After 48 hours, paraformaldehyde was added to each well to achieve a final concentration of 4% and the plate incubated for 1 hour at room temperature. The medium was removed, cells were stained with crystal violet (70% v/v H2O, 10% v/v ethanol, 20% v/v methanol and 0.25% crystal violet powder [Sigma Aldrich]) and washed three times with water. Cytopathic viral activity was determined by measuring absorbance of each well at 590 nm using a Varioskan LUX microplate reader. Drug activity was expressed as a percentage of inhibition of viral growth relative to the uninfected/untreated control (100% inhibition of viral cytopathic activity) and the infected/untreated control (0% inhibition of viral cytopathic activity) on that plate. Automated analysis was performed to maintain data integrity and objectively assess output. Non-linear regression was performed to generate concentration-effect predictions.

### In vivo studies

All work involving SARS-CoV-2 was performed at containment level 3 by staff equipped with respirator airstream units with filtered air supply. Prior to the start of the study, all risk assessments and standard operating procedures were approved by the University of Liverpool Biohazards Sub-Committee and the UK Health and Safety Executive. All animal studies were conducted in accordance with UK Home Office Animals Scientific Procedures Act (ASPA, 1986). Additionally, all studies were approved by the local University of Liverpool Animal Welfare and Ethical Review Body and performed under UK Home Office Project Licence PP4715265. Male Syrian golden hamsters (80-100 g; Janvier Labs) were housed in individually-ventilated cages with environmental enrichment under SPF barrier conditions and a 12-hour light/dark cycle at 21 °C ± 2 °C. Free access to food and water was provided at all times. Hamsters were randomly assigned into three groups of five and acclimatised for 7 days. Subsequently, hamsters (n=5 per group) were anaesthetised under 3% isoflurane and intranasally inoculated with either PBS (Group 1) or 100 ul of 1 × 10∧3 nCoV19 isolate SARS-CoV-2 Delta variant B.1.617.2 (Groups 2 and 3). Twenty-four hours post-infection (p.i) hamsters were treated, through intraperitoneal administration, with vehicle NaOH solution buffered to pH 7 (Groups 1 and 2). Group 3 were treated with probenecid in buffered NaOH solution at a dosage of 100 mg/kg. Treatment continued in a twice-daily dosing regimen for 48 hours p.i. On day 3 p.i all animals were ethically euthanised and lung lobes were harvested for quantification of viral load. In all cases, animal sacrifice was conducted via a lethal intraperitoneal injection of pentobarbitone, followed by cardiac puncture and immediate exsanguination of blood from the heart.

### Quantification of viral load by qPCR

A section of dissected lung lobes were homogenised in 1 mL of TRIzol reagent (ThermoFisher) using a bead mill homogeniser (Fisher Scientific) and Precellys CKmix lysing tubes (Bertin Instruments) at 3.5 metres per second for 30 seconds. The resulting lysate was centrifuged at 12,000 x g for 5 min at 4°C. Throat swab media (260 μL) was added to 750 μL of TRIzol LS reagent (ThermoFisher). The clear supernatants were transferred to Phasemaker™ tubes (ThermoFisher) and processed as per the manufacturer’s instructions to separate total RNA from the phenol-chloroform layer. Subsequently, the recovered RNA was precipitated using GlycoBlue™ according to the manufacturer’s instructions (ThermoFisher), washed and solubilised in RNAse-free water. The RNA was quantified and quality assessed using a Nanodrop (ThermoFisher). Samples were diluted to either 20,000 or 200 ng/mL in 60 μL of RNAse-free water. The resulting RNA samples were DNAse treated using the TURBO DNA-free™ kit according to the manufacturer’s instructions (ThermoFisher). The DNAse treated RNA was stored at −80°C prior to downstream analysis.

The viral RNA derived from hamster lung was quantified using a protocol adapted from the CDC 2019-Novel Coronavirus (2019-nCoV) Real-Time PCR Diagnostic Panel17 and a protocol for quantifying the SARS-CoV-2 subgenomic E gene RNA (E SgRNA)18 using the GoTaq^®^ Probe 1-Step RT-qPCR System (Promega). For quantification of SARS-CoV-2 using the nCoV assay, the N1 primer/probe mix from the SARS-CoV-2 (2019-nCoV) CDC qPCR Probe Assay (IDT) were selected. A standard curve was prepared (1,000,000 – 10 copies/reaction) via a 10-fold serial dilution of the CDC RUO 2019-nCoV_N_Positive Control (IDT). DNAse treated RNA at 200 ng/mL or dH2O was added to appropriate wells producing final reaction volumes of 20 μL. The prepared plates were run using a Chromo4™ Real-Time PCR Detector (Bio-Rad). The thermal cycling conditions for the qRT-PCR reactions were: 1 cycle of 45°C for 15 min, 1 cycle of 95°C for 2 min, followed by 45 cycles of 95°C for 3 seconds and 55°C for 30 seconds.

Quantification of SARS-CoV-2 E SgRNA was completed utilising primers and probes previously described elsewhere18 and were used at 400 nM and 200 nM, respectively (IDT), using the GoTaq^®^ Probe 1-Step RT-qPCR System (Promega). Quantification of 18S RNA utilised previously described primers and probe sequences19, and were used at 300 nM and 200 nM, respectively (IDT), using the GoTaq^®^ Probe 1-Step RT-qPCR System (Promega). Methods for the generatioplan of the 18S and E SgRNA standards have been outlined previously.20 Both PCR products were serially diluted to produce standard curves in the range of 5 × 108 - 5 copies/reaction via a 10-fold serial dilution. DNAse treated RNA at 20,000 ng/mL or dH2O were added to appropriate wells producing final reaction volumes of 20 μL. The prepared plates were run using a Chromo4™ Real-Time PCR Detector (Bio-Rad). The thermal cycling conditions for the qRT-PCR reactions were: 1 cycle of 45°C for 15 min, 1 cycle of 95°C for 2 min, followed by 45 cycles of 95°C for 3 seconds and 60°C for 30 seconds. Both N and E SgRNA data were normalised to 18S data for subsequent quantitation.

### Quantification of viral load by plaque assay

Vero E6 plaque assays were performed for quantification of plaque formation within individual samples. A section of dissected lung lobes were placed in screw-top microcentrifuge tubes containing a single stainless steel bead cooled to 4 °C. 500 μL EMEM (Gibco; 670086) was added to each microcentrifuge tube and the lung tissue homogenised using a TissueLyser LT (Qiagen, 85600) for approximately 4-5 minutes at 50 Hz. Microcentrifuge tubes were then centrifuged at 2,000 rpm for 5 minutes at room temperature. The homogenised tissue supernatant was then collected, and stored at −80 °C. Homogenised samples were thawed, diluted in EMEM (1:4 1:20, 1:100, 1:500, 1:2500, and 1:12,5000), and layered over confluent Vero E6 cells in 100 μL volumes, in triplicate, in 96-well plates. 100 μL semi-solid media was then added to each well. The plates were then incubated at 37°C with 5% CO2. After 72 hours, paraformaldehyde was added to each well to achieve a final concentration of 4% and the plate incubated for 1 hour at room temperature. The medium was removed, cells were stained with crystal violet and washed three times with water. The number of plaques in each well were enumerated at the highest countable concentration. The average value was used to calculate the concentration of each sample in viral plaque forming units (PFU).

### Statistical analysis

An unpaired t-test was used to compare differences in body weight between probenecid treated and vehicle treated groups on day 3 post infection using R (V.4.2.1) (24).

## Results

### In vitro Vero E6 cell assay

7-point concentration-response analysis was performed in triplicate with three independent biological replicates. All plates passed quality control. The control compound, remdesivir, generated a robust 4-parameter fit (Figure 1) – EC50 = 2.43 μM, EC90 = 9.39 μM, EMAX = 98.86 and hillslope = 1.80. No detectable activity was observed for probenecid at any concentration up to 25 μM.

**Figure 1.**
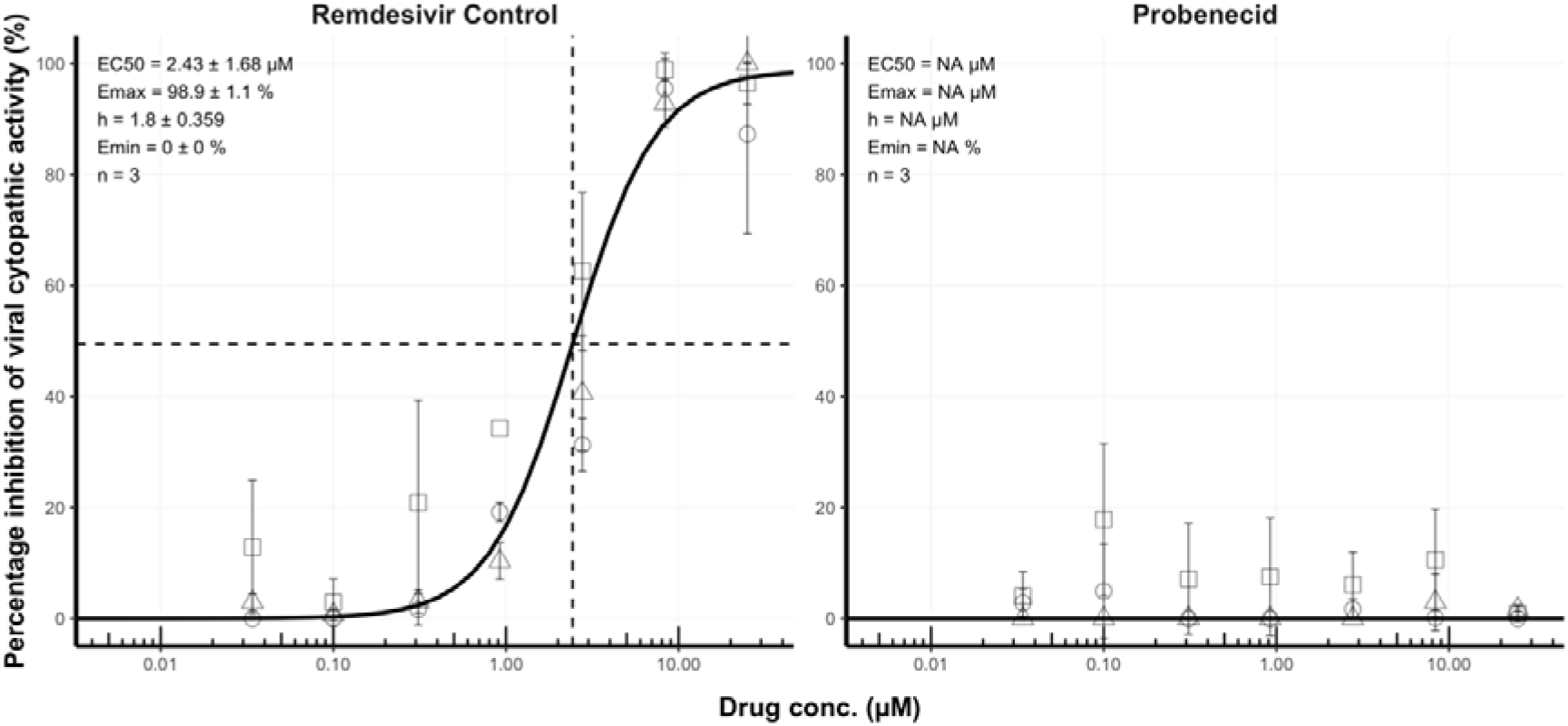
Concentration-effect relationship for the inhibition (%) of SARS-CoV-2 cytopathic activity for remdesivir and probencid. For each compound, activity was expressed relative to uninfected/untreated controls (100% inhibition of viral cytopathic activity) and infected/untreated controls (0% inhibition of viral activity). For each compound, we assessed activity at 25.00 μM, 8.33 μM, 2.78 μM, 0.93 μM, 0.31 μM, 0.10 μM and 0.03 μM in triplicate. Non-linear regression using an EMAX model was performed on data taken from three independent biological replicates to generate concentration-effect predictions (solid black lines). EC50 values, hillslope and replicate number (n) are shown. Dashed lines represent the EC50. Squares, diamonds and circles represent individual biological replicates and error bars represent standard deviation calculated from technical triplicates.

**Table 1.** Differences between previous and current experimental approaches for a) in vitro analysis and b) in vivo investigations.

|  |  |  |
| --- | --- | --- |
| **a** | **Murray *et al.*** | **Jeffrys *et al.*** |
| **Duration of exposure p.i. (hr)** | 96 | 48 |
| **MOI** | 0.01 | 0.05 |
| **Detection method** | Plaques/ well | Percentage cytopathic activity<br>relative to control |
| **b** | **Murray *et al.*** | **Box *et al.*** |
| **Dose** | 200 mg/kg OD | 100 mg/kg BID |
| **Virological Outcome Measures** | PFU/mL | Copies/mL (total and SgE) |
| **Strain** | B16 | B.1.617.2 (Delta) |

### In vivo hamster model of SARS-CoV-2 infection

Hamsters were inoculated with virus and at 24 hpi were treated with probenecid, (i.p. 100 mg/kg BID) for 4 doses. Hamster weight was monitored throughout the study as a marker for health. Figure 2 shows animal weight relative to baseline (day 0; prior to SARS-CoV-2 inoculation). All animals displayed moderate weight loss 24 hours following infection (4-7% of bodyweight) regardless of treatment. Mean body weight remained relatively consistent in uninfected animals throughout the study (Figure 2). To determine the viral load in animals infected with SARS-CoV-2 and dosed with either the vehicle control or probenecid, total RNA was extracted from the lung samples harvested on day 3 post infection. Viral replication was quantified using qRT-PCR to measure total and sub-genomic viral RNA relative to the E gene (sgE) as a proxy. These data are illustrated in Figure 3. There is no apparent reduction in either total lung or sgE RNA for probenecid treated animals compared to infected controls (P = >0.5). RNA levels for uninfected control samples were below the assay lower limit of quantification (Figure 3).

**Figure 2.**
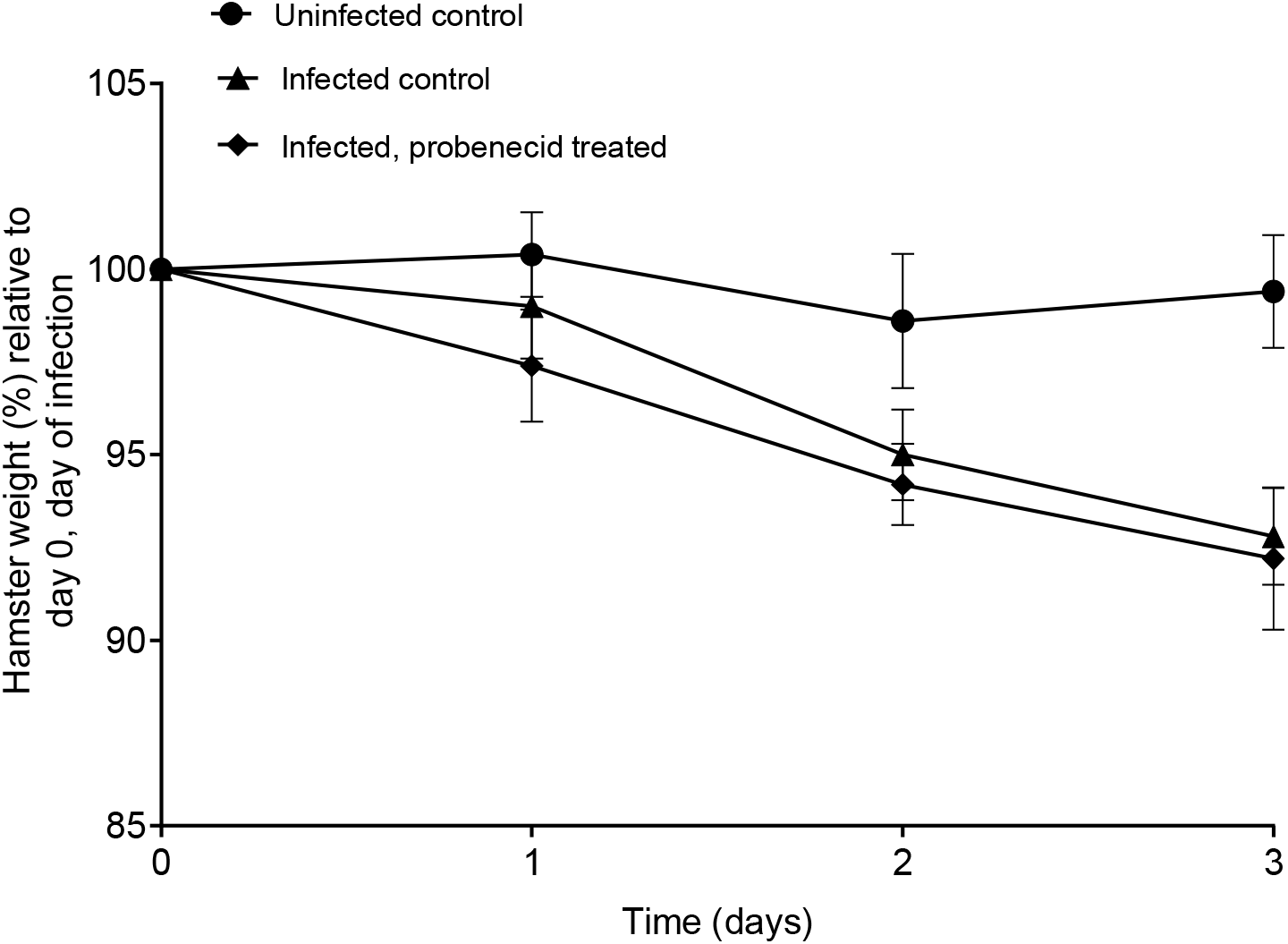
Average body weight data post-infection. Error bars represent the standard deviation between individual animal weights.

**Figure 3.**
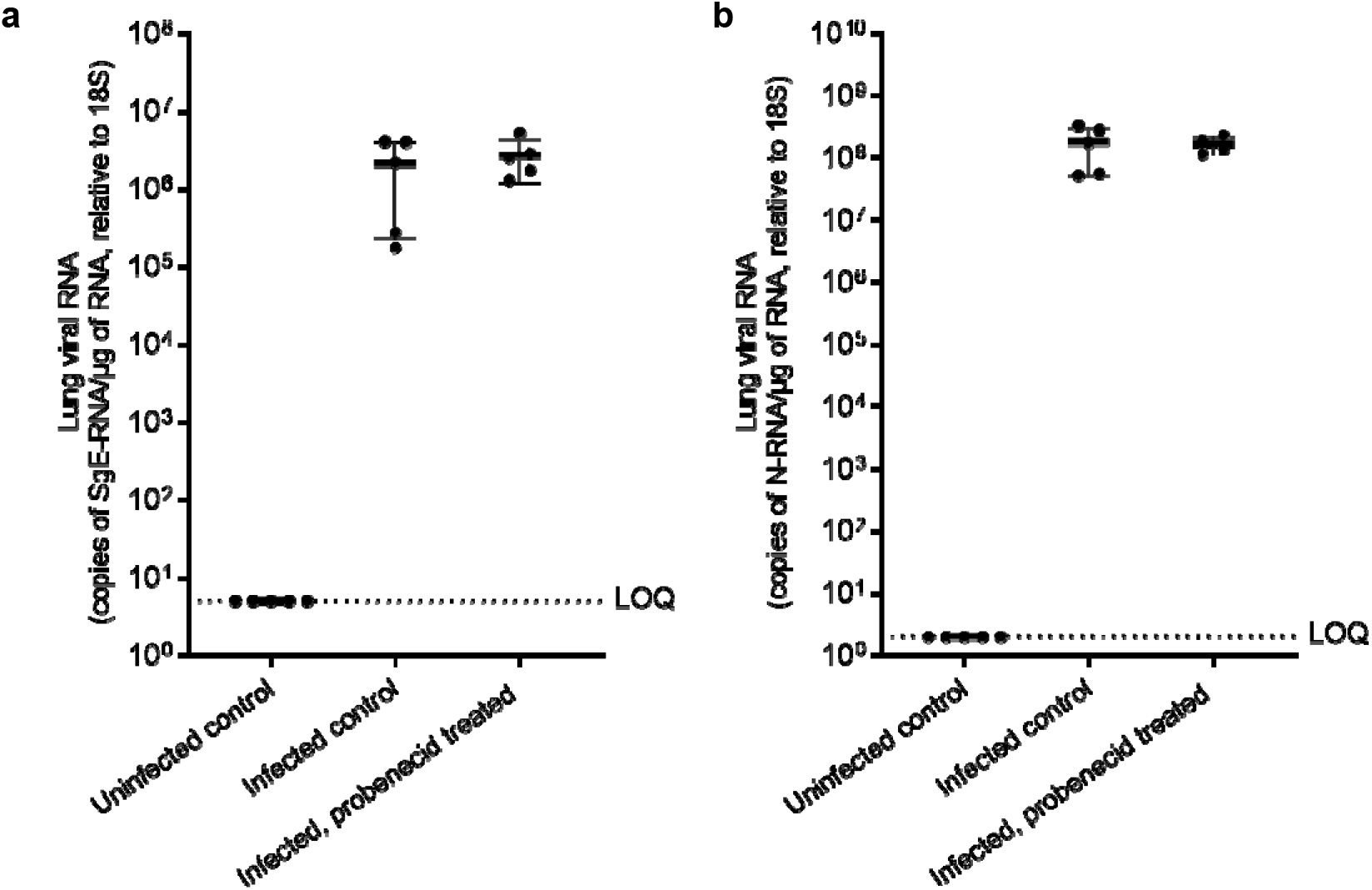
Lung viral RNA normalised to the 18S subunit (a) and total RNA (b) in untreated controls and SARS-CoV-2 infected animals, treated with vehicle or probenecid. Error bars represent standard deviation between samples obtained from individual animals.

**Figure 4.**
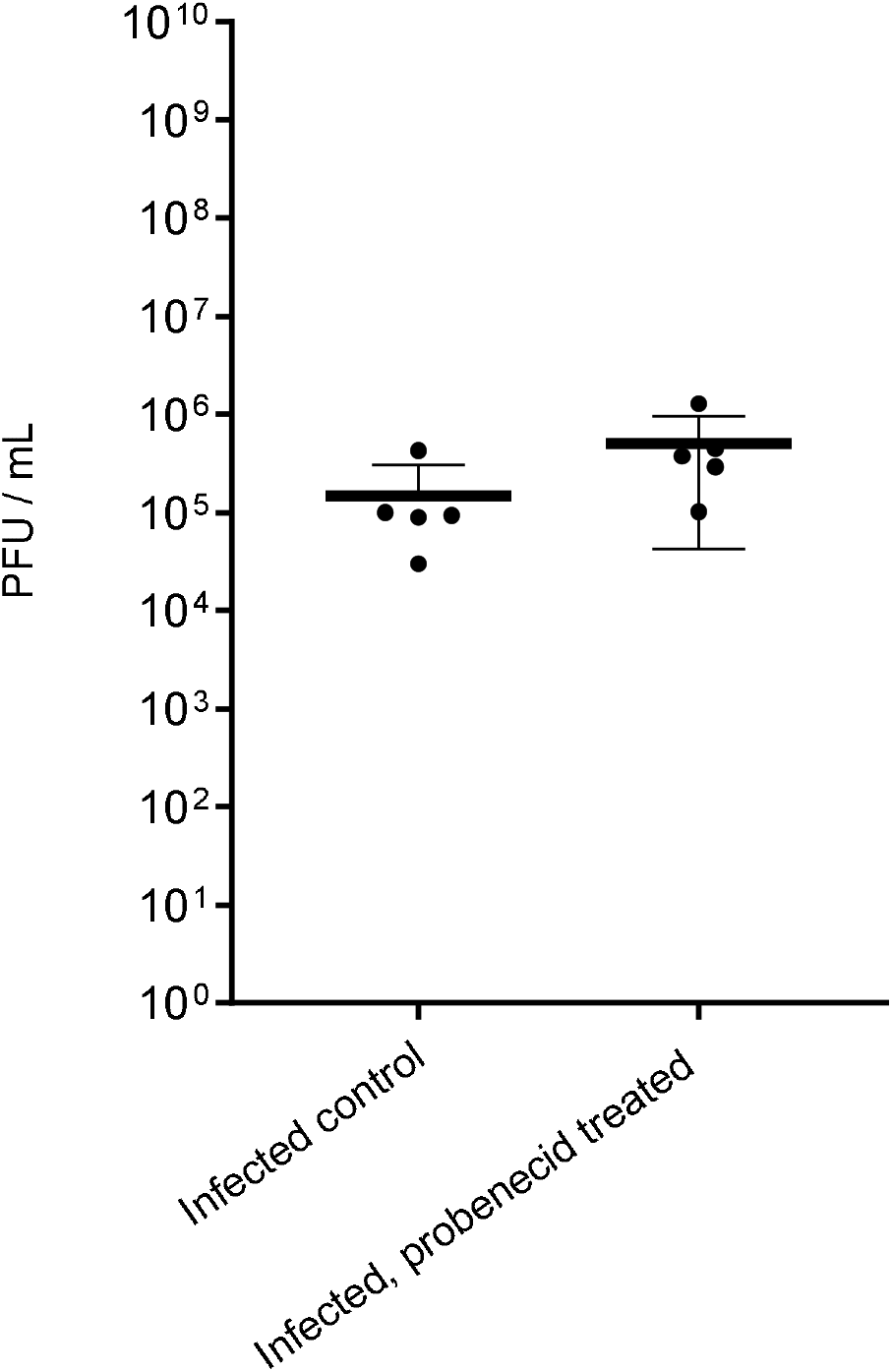
Plaque forming units (PFU/mL) in terminal lung samples from SARS-CoV-2 infected animals, treated with vehicle (n=5) or probenecid (n=5). Error bars represent standard deviation of values obtained for individual animals.

## Discussion

Re-purposed agents usually exert their activity via fortuitous similarity in a target or via a secondary mechanism of action which is often poorly understood. Accordingly, a repurposed drug cannot necessarily be expected to exhibit levels of potency that can be achieved through development of a mechanism-based inhibitor. Consequently, it is expected that much more potent antiviral drugs for SARS-CoV-2 will emerge in the months and years to come. Nonetheless, drugs such as molnupiravir and nirmatrelvir have shown early evidence of efficacy in COVID-19 demonstrating the utility of repurposing drugs in an emergency pandemic situation. For drugs with sub-optimal potency the use of drug combinations accessing complimentary mechanisms of action should be considered as an urgent short-term research priority to maximise efficacy and mitigate the resistance risk, which are currently understudied for existing antiviral drugs being deployed for COVID-19.

The speed at which preclinical methodologies have been developed in the first years of the pandemic is laudable but substantive inter-laboratory differences in assay conditions and outcomes are increasingly evident. However, the presented data do not support probenecid for treatment of COVID-19 and indicate that probenecid does not significantly inhibit SARS-CoV-2 infection at dosages where efficacy was observed elsewhere (22). Notable differences in methodology between the current study and the previous report, render it difficult to discern with certainty which of the outcomes is most representative of the ultimate candidacy of probenecid. It should be noted, however, that animal data to date are largely representative of treatment models for severe disease where antivirals may also be deployed in pre- or post- exposure prophylaxis, mild to moderate disease.

Plasma and tissue-site protein binding is known to impact the likelihood of success for some, but not all, drug therapies. Protein binding should therefore be carefully considered when interpreting candidacy of a putative antiviral intervention (25). This approach would ensure that PK-PD targets are clearly defined, and that unbound drug concentrations at the target site can reasonably be expected to remain above the efficacy threshold for the duration of the dosing interval (26). It is important to note that proposed target CMAX / EC90 ratios for probenecid against SARS-CoV-2 in humans are yet to be investigated and that neither the current study, nor the previous study, empirically investigated the consequences of protein binding directly. Murray et al. (2021) did correct the in vitro-derived EC_90_ value for 95% plasma protein binding when deriving the target for their pharmacokinetic simulations, but the absence of empirically determined protein-adjusted EC90 values prevents a robust assessment of it’s importance for the application of probenecid.

Notwithstanding, Murray et al. demonstrated that target exposures could reach the reported EC90 values, despite a highly stringent correction for protein binding.

There are several factors that may have contributed towards the differences in outcome between the current study and that of Murray et al. OAT3 (SLC22) has been suggested as the host antiviral target for probencid, as previously reported for influenza A (21). However, the role of OAT3 in SARS-CoV-2 replication has not been empirically investigated and neither have potential differences in the OAT3 expression in the different cell lines employed. Whatever the putative mechanism of action for probenecid, differences between different cell systems and culture conditions in different laboratories cannot be ruled out. It is assumed that, as with other antiviral agents, the pharmacodynamic indices, relative to efficacy, are concentration-dependent but no data for SARS-CoV-2 are currently available (27). Depending on the interaction of probenecid to its target, it is not possible to determine whether efficacy may be driven in a concentration- or time-dependent manner (e.g time > EC90, AUC/ EC90 or, Cmax/ EC90). Differences in the duration of in vitro experiments (48 vs 96 hours) may therefore have influenced the observed differences in outcome. However, this would not explain the observed differences in virological outcome seen in the animal experiments since no efficacy was seen in the current study despite multiple dosing of an absolute dose that was shown to be effective by Murray et al.

Re-purposed antiviral agents need to be safe, affordable, readily available for scale-up and production for clinical investigations and preclinical data need to be robust and reproducible between laboratories. There is a critical need to better understand the impact of assay conditions so that methodology can be refined to provide definitive recommendations. Data presented here highlights the significance of differences in preclinical methodology and the importance of understanding inter-laboratory variation to provide an evidence-based decision to inform candidate selection for clinical trials.

## Acknowledgments

The authors thank Ralph Tripp and Jackelyn Crabtree at the University of Georgia for highly constructive discussions to understand differences in methodology.

